# Single nucleotide polymorphisms shed light on correlations between environmental vairalbe and adaptive genetic divergence in Chum salmon (*Oncorhynchus keta*)

**DOI:** 10.1101/001974

**Authors:** Xilin Deng, Philippe Henry

## Abstract

Identifying the genetic and ecological basis of adaptation is of immense importance in evolutionary biology. In our study, we applied a panel of 58 biallelic single nucleotide polymorphisms (SNPs) for the economically and culturally important salmonid *Oncorhynchus keta.* Samples included 4164 individuals from 43 populations ranging from Coastal Western Alaska to southern British Colombia and northern Washington. Signatures of natural selection were detected by identifying seven outlier loci using two independent approaches: one based on outlier detection and another based on environmental correlations. Evidence of divergent selection at two candidate SNP loci, *Oke_RFC2-168 and Oke_MARCKS-362*, indicates significant environmental correlations, particularly with the number of frost-free days (NFFD). Important associations found between environmental variables and outlier loci indicate that those environmental variables could be the major driving forces of allele frequency divergence at the candidate loci. NFFD, in particular, may play an important adaptive role in shaping genetic variation in *O. keta.* Correlations between divergent selection and local environmental variables will help shed light on processes of natural selection and molecular adaptation to local environmental conditions.

## Introduction

Chum salmon, *Oncorhynchus keta*, has the largest range among all Pacific salmon species in the North Pacific Rim (Sato *et al*. 2004), and it constitutes an important part of the Pacific Rim ecosystem (Seeb *et al*. 2011). All Pacific salmon species are anadromous with the exception of some species, particularly sockeye and masu salmom, having nonanadromous populations (Quinn 2005). They hatch in fresh water, then migrate to sea where they spend most of their lives, and return to fresh water to spawn at maturity. Pacific salmon have a strong tendency to return from ocean feeding areas to their natal streams to spawn, and this strong homing behavior can result in reproductive isolation of populations (Quinn 2005).

Pacific salmon vary greatly in age and size at maturity, morphology, and timing of spawning (Beacham and Murray 1987). Differences in spawning time and location result in distinct salmon stocks. Stocks might be genetically different due to their adaptations to their particular spawning locations; different spawning, incubation, and rearing environments may thus cause their adaptive genetic differences (Beacham and Murray 1987).

Patterns of environmental variation can shape adaptive genetic variation across a species’ range. Environmental heterogeneity subjects populations to varying selective pressures, and this may lead to local adaptation (Schoville *et al*. 2012). Natural selection can play a major diversifying role in salmonid populations, and environmental variables such as water temperature, stream size, female choice, and predation risk have been found to be among the most influential agents of natural selection in this system (Garcia de Leaniz *et al*. 2007). Thus, correlations between genetic variation and environmental gradients can be interpreted as evidence of natural selection by uncovering loci that are linked to selected genes (Eckert *et al*. 2010).

Genome-wide scans of patterns of single nucleotide polymorphisms (SNPs) can reveal patterns of adaptive genetic variation by detecting locus-specific signatures of positive selection. An alternative way of uncovering signs of local adaptation is by identifying significant correlations between genetic polymorphisms and environmental variables (Coop *et al*. 2010). Nonetheless, associations found between environmental variables and adaptive genetic divergence do not necessarily indicate causal relationships, but they can provide insights into the selective forces potentially generated by environmental variables in natural selection (McColl and McKechnie, 1999).

The purpose of this study was to detect signatures of local adaptation in *O. keta* by investigating the correlations between patterns of allele frequency differentiation and environmental gradients. We hypothesize that given the wide range of environmental conditions encountered by *O. keta*, there are corresponding adaptive genetic differences among the salmon populations. We apply a panel of 58 SNPs to 43 populations of chum salmon ranging from coastal western Alaska to southern British Colombia and northern Washington. We apply the SNP database to detect signatures of natural selection in *O. keta* by identifying SNP outlier loci in terms of the genetic differentiation index *F*_*ST*_. In addition, we examine the correlations between allele frequencies at outlier loci and environmental variables.

## Methods

### Data collection

In the present study, we obtained a published data set (Seeb *et al*. 2011) available from the online repository DRYAD (www.datadryad.org). We selected a subset of the representative populations distributed throughout the Pacific Northwest (Fig 1). As a result, a total of 58 biallelic single nucleotide polymorphisms (SNPs) genotyped in 4164 individuals from 43 different populations of chum salmon were studied. The data set was sampled in regions ranging from Coastal Western Alaska to Southern British Colombia and Northern Washington (Fig. 1). Additionally, we collected sixty monthly, seasonal and annual environmental variables related to temperature, precipitation and topography for each sampling location using climateWNA (http://www.genetics.forestry.ubc.ca/cfcg/ClimateWNA/ClimateWNA.html; Wang *et al*. 2012).

**Figure 1.**
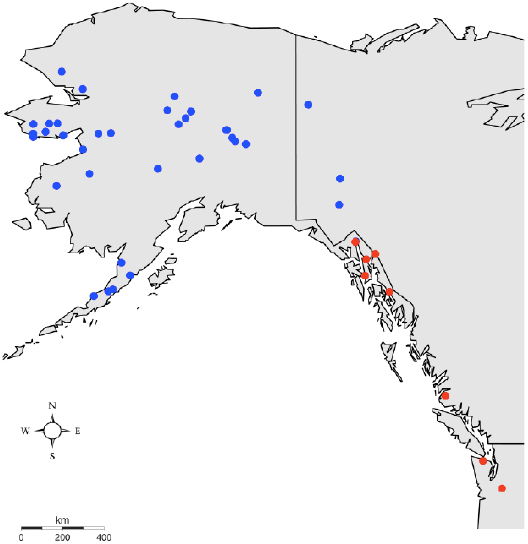
Map of sampling locations for 43 populations of chum salmon. Blue symbols indicate populations from the northern lineage; red symbols indicate populations from the southern lineage as identified using STRUCTURE.

### Genetic variation

All SNP markers have already been tested to conform to Hardy-Weinberg (HW) equilibrium in Seeb *et al*. (2011), and four linked SNP pairs were found to display patterns of significant gametic disequilibrium. Since linkage was only observed within a handful of populations, we retained all SNP markers for further analyses.

### Environmental variables and PCA

A principal component analysis (PCA) using the statistical software R (package ade 4) was applied to all two hundred and sixteen environmental variables to examine possible correlations between all variables conducted. Variables that were correlated at |r| > 0.8 were considered redundant and were thus removed (Manel 2010). Within each pair of variables that were highly correlated, we kept only one biologically relevant variable. Therefore, only environmental variables identified as being uncorrelated by the PCA analysis were used.

### Population structure

The extent of population differentiation was quantified by calculating pairwise *F*_*ST*_ for each locus and over all loci among all 43 chum salmon populations using Arlequin 3.5 (Excoffier *et al*. 2009). In addition, we used the software STRUCTURE 3.4 (Pritchard *et al*. 2000) to detect clusters (K) of individuals. We ran 10 chains of 100,000 iterations with K ranging from 1 to 10. We then ran the structure program with the exclusion of the outlier loci to investigate if the outliers had an effect on the population structure. We used STRUCTURE HARVESTER Web v0.6.92 (Earl and vanHoldt 2012) to estimate Delta *K* (Δ*K*) and (ln *P*(*K*)), the natural log probability of K.

### Signature of natural selection

Arlequin 3.5 (Excoffier *et al*. 2009) was used to detect outlier loci. We ran 100,000 simulations assuming 100 demes per group using a finite island model without taking into account the underlying population structure. We then ran the software again using the hierarchical island model by grouping populations into two groups representing the northern and the southern lineages based on the population clusters identified by STRUCTURE. Loci that fell above the 99% quantile were determined to be under positive selection, and loci that fell below the 99% quantile were considered to be candidates for balancing selection.

### Environmental effects on adaptive genetic variation

In order to reinforce evidence of natural selection acting on outlier loci, we applied an alternative approach that utilizes associations between environmental variables and patterns of allele frequencies in identifying genetic adaptive variation. Allele frequencies are typically correlated among geographically closer populations; as a result, the association found between environmental variables and allele frequencies could occur by chance due to isolation by distance or similar environmental factors acting on geographically proximate populations (Limborg *et al*. 2011; Coop *et al*. 2010). If not taken in to account, signals from neutral population structure could lead to a high false positive rate (Coop *et al*. 2010). To overcome false positivity caused by neutral population structure, we accounted for spatial autocorrelation when testing for correlations between allele frequencies and environmental variables. Applying 150,000 iterations in Bayenv (Coop *et al*. 2010), we tested for correlations between the SNP allele frequencies and the following variables that were identified as being uncorrelated by the PCA analysis: (1) precipitation as snow (mm) (PAS), (2) degree-days above 5°C, growing degree-days (DD5), (3) mean annual precipitation (mm) (MAP), (4) the number of frost-free days (NFFD), (5) degree-days above 18°C, cooling degree-days (DD_18), (6) mean warmest month temperature (°C) (MWMT), (7) annual heat:moisture index (MAT (mean annual temperature (°C))+10)/(MAP/1000)) (AHM), (8) temperature difference between MWMT and MCMT (mean coldest month temperature (°C), or continentality (°C) (TD), (9) the Julian date on which FFP begins (bFFP), (10) mean annual summer (May to Sept.) precipitation (mm) (MSP), (11) summer heat:moisture index ((MWMT)/(MSP/1000)) (SHM), (12) longitude, (13) latitude and (14) elevation.

## Results

### Genetic diversity

Among all chum salmon populations, varying levels of differentiation were revealed with values ranging from 0.0311 to 0.157 and the average over all SNP markers was 0.0564. Genetic diversity was measured by average heterozygosity per locus and the level of genetic diversity ranged from 0.224 to 0.299 (He), 0.227 to 0.304 (Ho) in all 43 populations (Table 1).

**Table 1.**
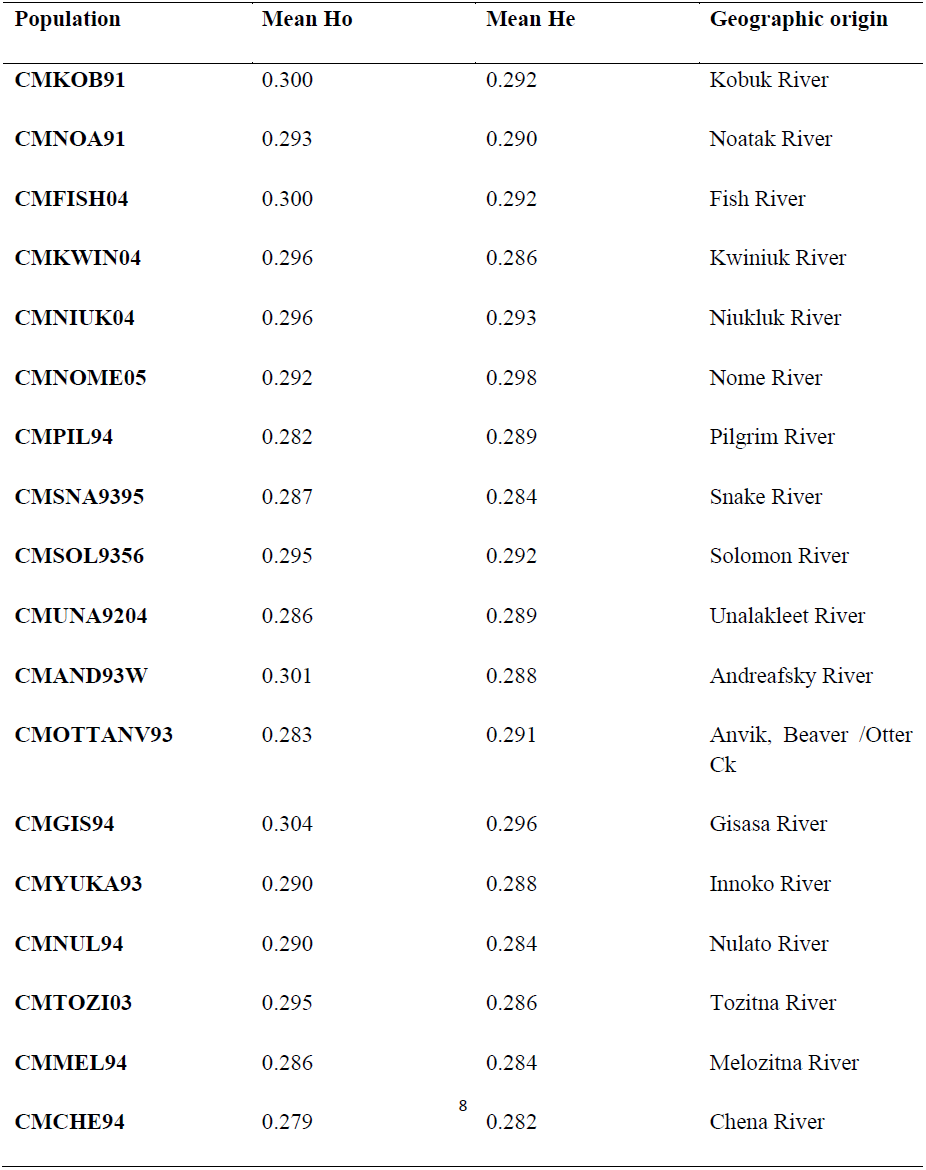
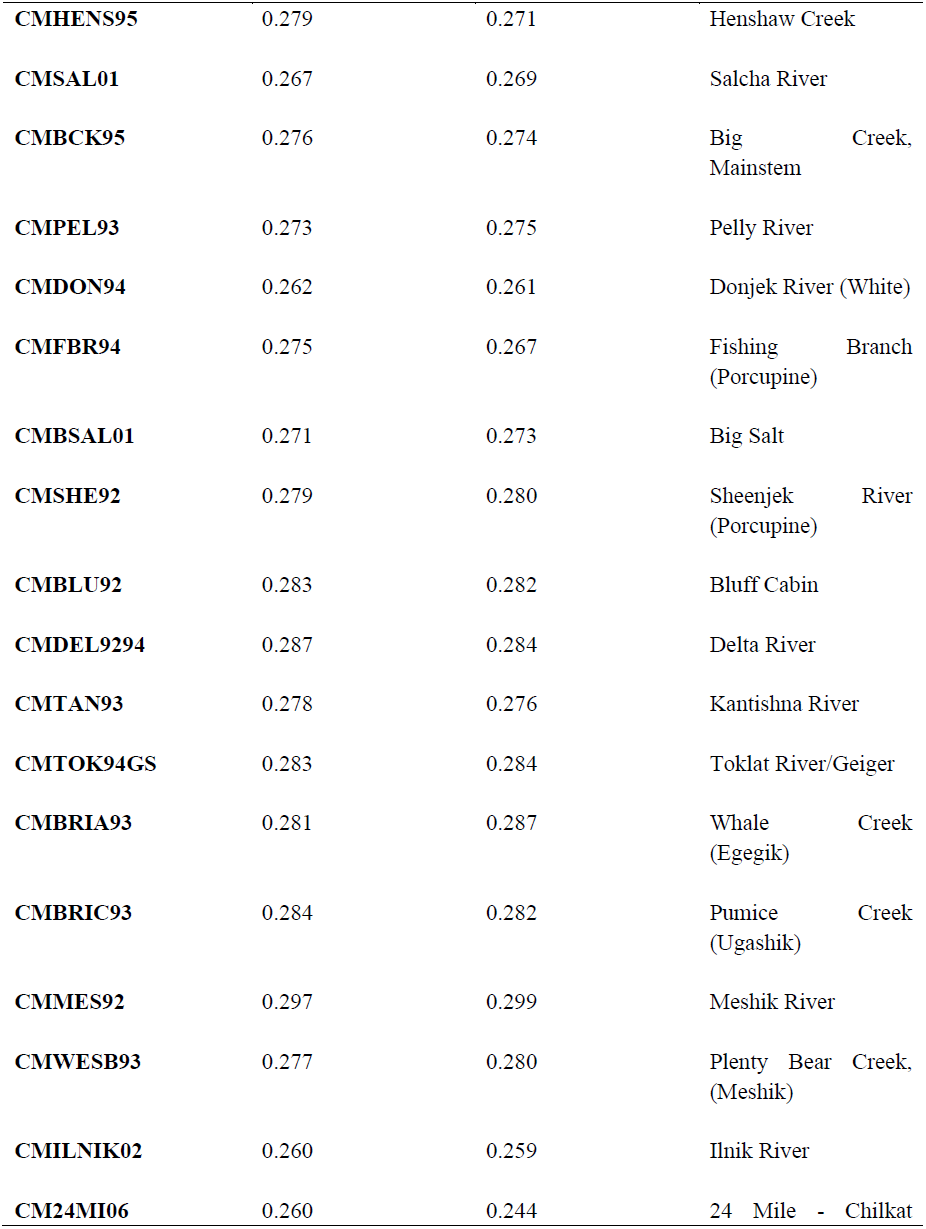
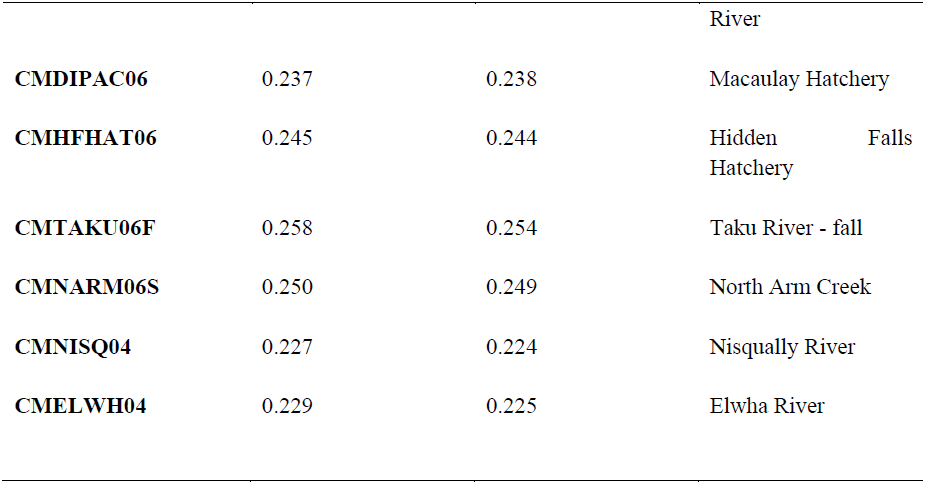
Location of spawning populations, average observed heterozygosity (Ho) and average expected heterozygosity (Seeb *et al*. 2011).

### PCA analysis of environmental variables

Fourteen environmental (climatic and topographic) variables were identified as being uncorrelated from the PCA analysis and were therefore retained (Fig. 2)

**Figure 2.**
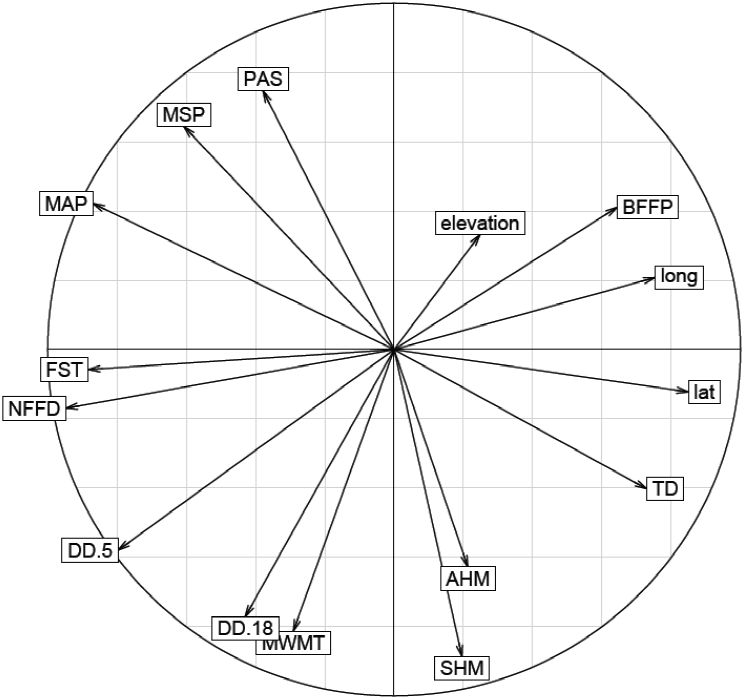
Representation of the fourteen retained environmental (climatic and topographic) variables on a principle component plot.

### Detection of loci under selection

The genome scan assuming a finite island model showed an excess of outlier loci (Fig. 3a). Undetected population substructure can often have negative effects on outlier tests for SNP loci under selection by causing inflated false-positive rates (Huelsenbeck and Andolfatto 2007), we thereby took into account the hierarchical population structure when running the Arlenquin test. Outlier loci were thus significantly reduced after accounting for a hierarchical structure based on the population clusters (K=2) previously identified by STRUCTURE. Using the hierarchical island model, a total of six outlier loci were detected with Arlenquin. We revealed three significant outliers for divergent selection (P<0.01) at the *F*_*ST*_ level (Fig. 3b), *Oke_FARSLA-242, Oke_u1–519* and *Oke_RFC2-168*, of which two loci were also candidates at the *F*_*CT*_ level, *Oke_u1–519* and *Oke_RFC2-168* (Fig. 3c). Outlier, *Oke_MARCKS-362*, was detected only at the *F*_*CT*_ level as a potential candidate for divergent selection. Two loci that lie below the 99% quantile were considered as candidates under balancing selection, *Oke_TCP1-78* and *Oke_arf-319* (Fig. 3b and 3c). Bayenv confirmed two out of the six outliers (p < 0.01) found with Arlenquin, *Oke_RFC2-168* and *Oke_MARCKS-362*, and detected a new outlier, *Oke_Tf-278*.

**Figure 3.**
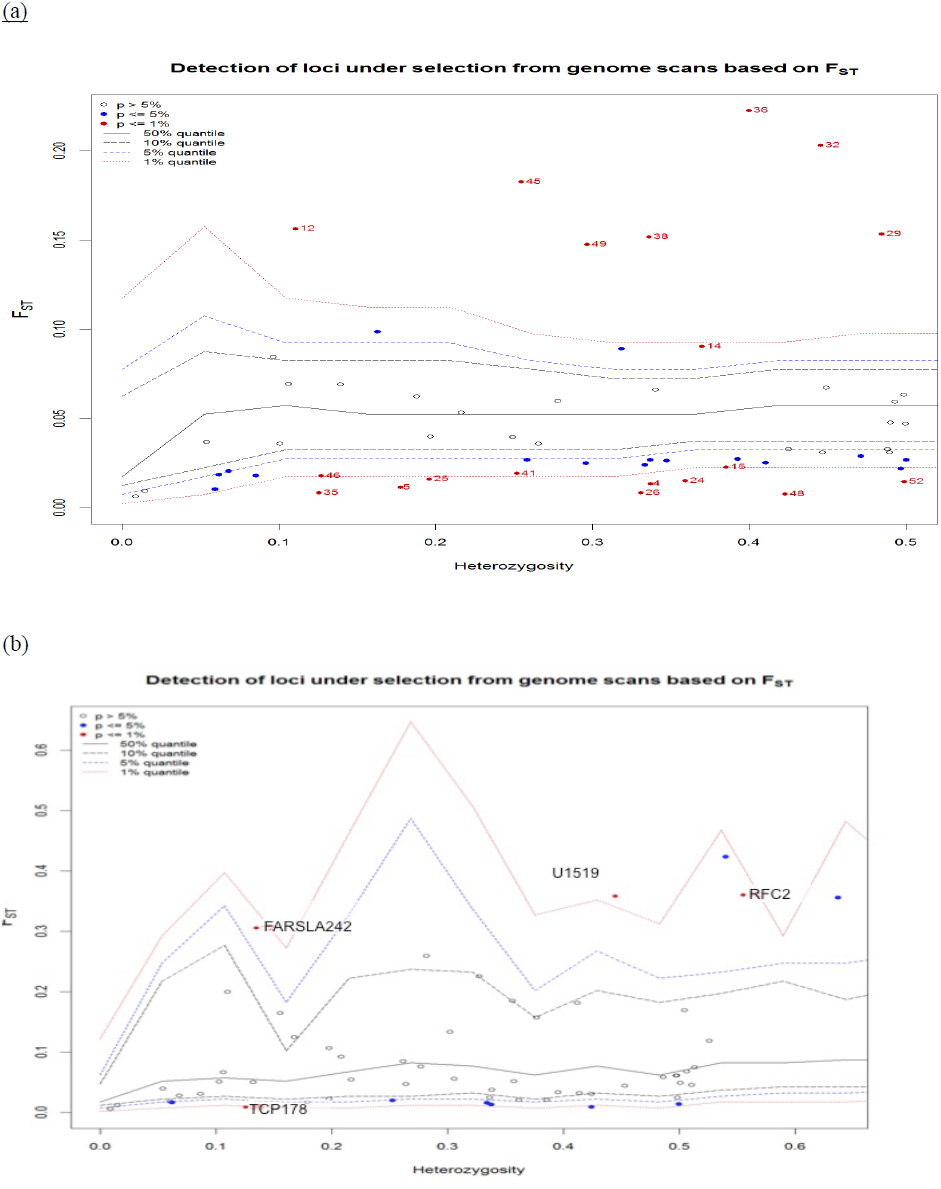
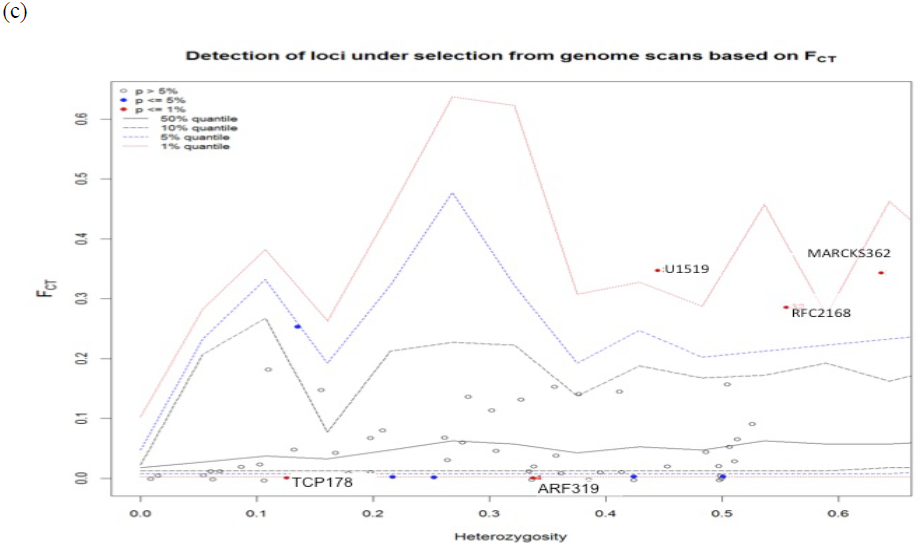
Outlier tests for detection of loci under selection using the method of *Excoffier et al.* (2009). (a) *F*_*ST*_-based test plotted against heterozygosity under the assumptions of a finite island model. (b) *F*_*ST*_-based test assuming a hierarchical island model by grouping all populations into two major groups. (c) *F*_*CT*_-based test assuming a hierarchical island model as in (b). The red solid lines represent the 1% quantiles from coalescent simulations; dashed lines indicate the 5% and 10% quantiles.

### Environmental correlates

Allele frequencies at the two SNP loci identified by Arlequin as being under positive selection (P<0.01)*, Oke_RFC2-168 and Oke_MARCKS-362*, were significantly correlated with one or two variables (Table 2). One new outlier, *Oke_Tf-278,* detected by Bayenv was found to associate with one environmental variable, the number of frost-free days (NFFD) (Table 2). The rest of the outlier loci, *Oke_FARSLA-242*, *Oke_TCP1-78*, *Oke_u1–519* and *Oke_arf-319*, were not correlated with any environmental variable(s) by Bayenv.

**Table 2.**
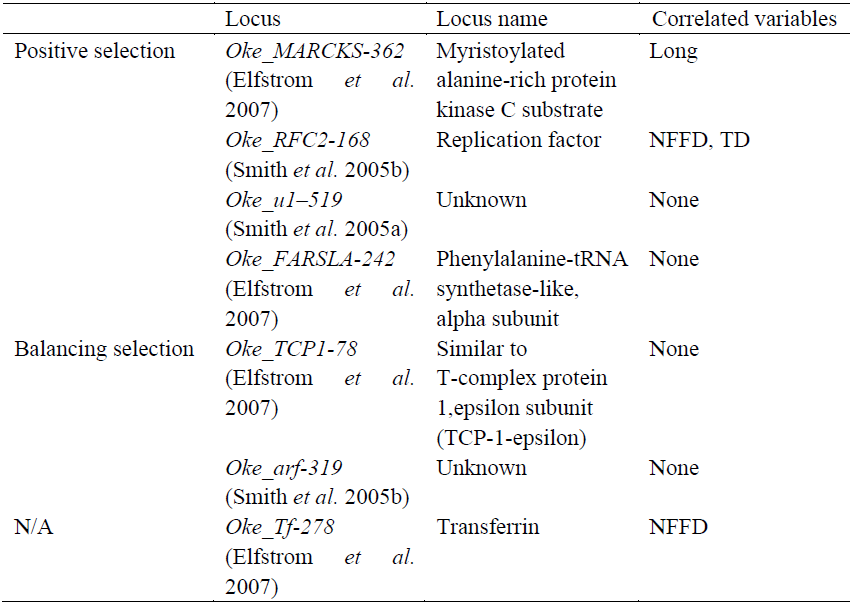
Summary of seven outlier loci, their locus names and results from Bayesian inference for correlation between allele frequencies and environmental variables.

Bayenv method revealed a strong correlation between allele frequencies at *Oke_RFC2-168* and the number of frost-free days (NFFD). As the number of frost-free days increases, the allele frequency decreases (Fig. 4). When the number of frost-free days (NFFD) reaches beyond 200 days, the allele frequency at *Oke_RFC2-168* tends towards a value near zero. One dot indicating a northern population stands apart due to its geographic proximity to the southern populations, hence similar number of frost-free days. However, it maintains its higher allele frequency due to its northern lineage.

**Figure 4.**
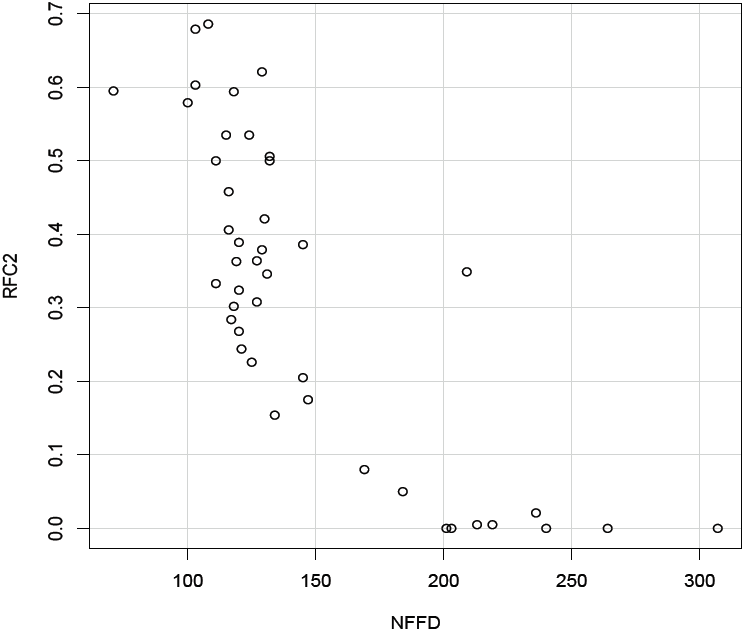
The correlation between allele frequencies at locus *RFC2-168* and the number of frost-free days (NFFD). Frequency of only a single allele is given for the SNP locus.

The allele frequency at *Oke_RFC2-168* is also positively correlated with TD (the temperature difference between the mean warmest month temperature (°C) and the mean coldest month temperature (°C)). The allele frequency increases steadily from 0 to 0.7 as TD increases from 0 °C to 45°C (Fig. 5).

**Figure 5.**
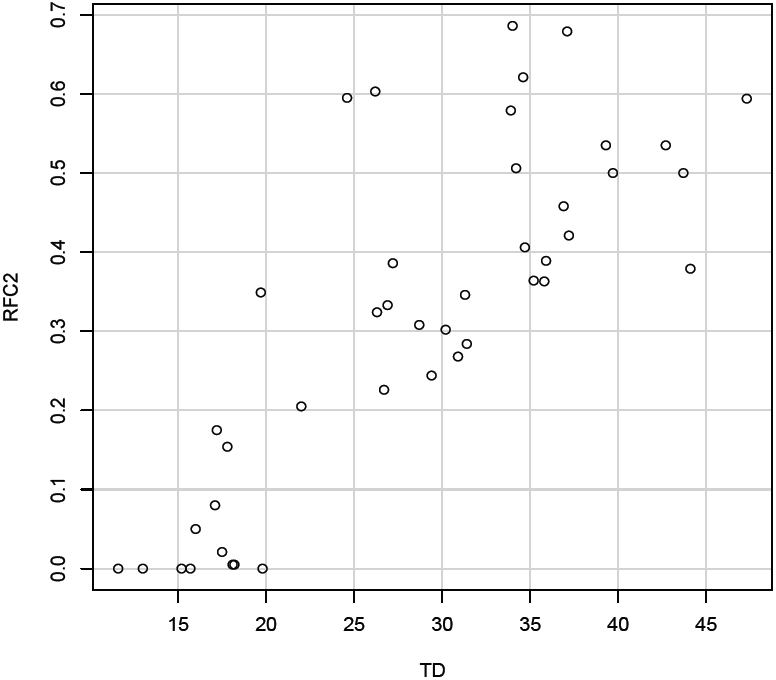
The correlation between allele frequencies at *Oke_RFC2-168* and TD (the temperature difference between the mean warmest month temperature (°C) and the mean coldest month temperature (°C)). Frequency of only a single allele is given for the SNP locus.

The allele frequency at *Oke_MARCKS-362* varied with longitude. The allele frequency is relatively high when longitude is less than 140 degrees (southern populations), and it decreases substantially and stabilizes after longitude increases beyond 140 degrees (northern populations) (Fig. 6).

**Figure 6.**
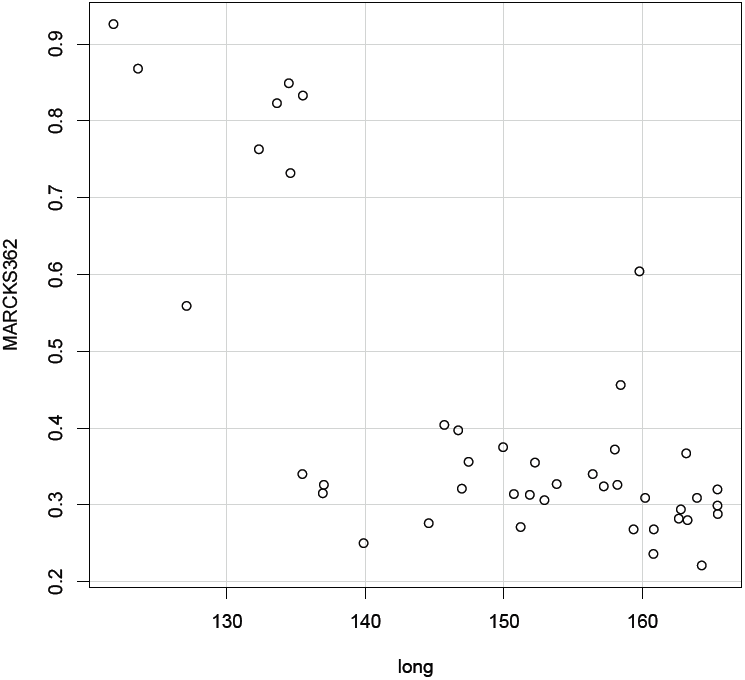
The correlation between allele frequencies at *Oke_MARCKS-362* and longitude (Long). Frequency of only a single allele is given for the SNP locus.

The allele frequency at *Oke_Tf-278* is positively correlated to NFFD and it increases as NFFD increases from 0 to 200 days (Fig. 7).

**Figure 7.**
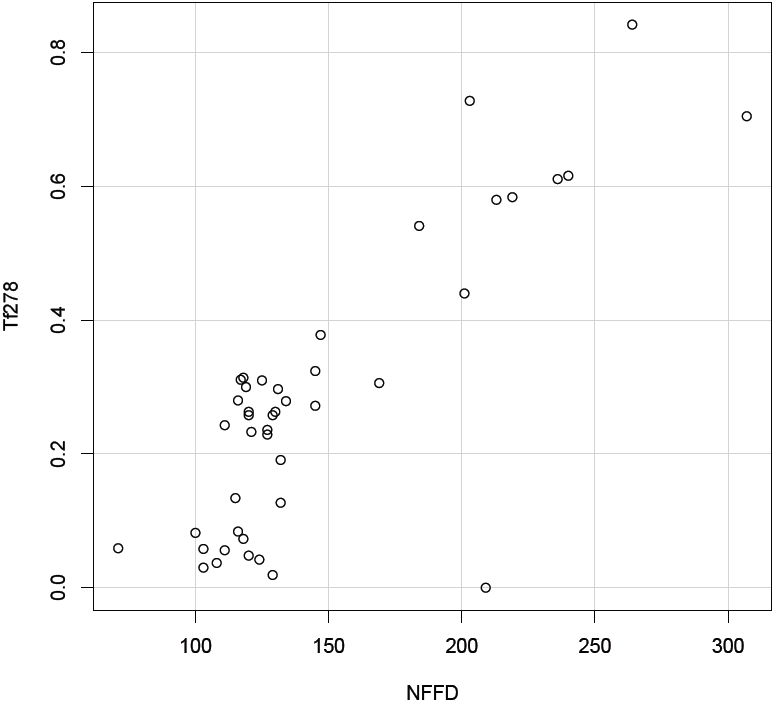
The correlation between allele frequencies at *Oke_Tf278* and the number of frost-free days (NFFD). Frequency of only a single allele is given for the SNP locus.

### Population structure

A strong neutral population structure was revealed (Fig. 8). Cluster analysis with STRUCTURE grouped populations into K=2 clusters. There are clear distinctions between two population clusters (Fig. 8a); after all six outliers found with Arlenquin were removed, however, the distinction between the two clusters was blurred (Fig. 8b).

**Figure 8.**
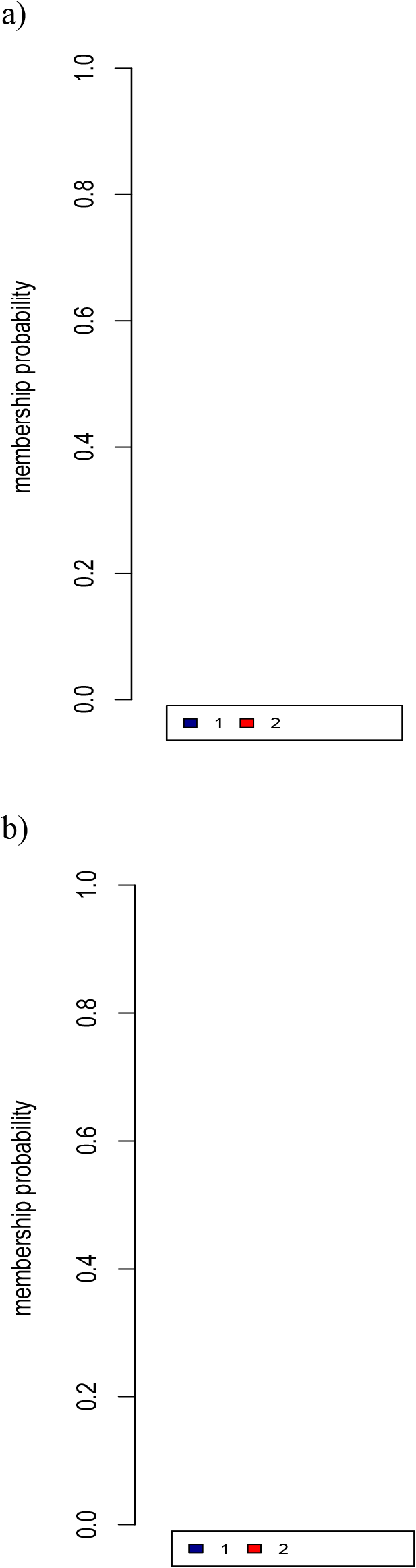
Number of population clusters (K) detected by STRUCTURE program 2.0 (a) test run by including all 58 SNP loci, and (b) test run by excluding all six outliers detected by Arlenquin.

## Discussion

In our study, significant associations between environmental variables and several outlier loci were found. These loci are potentially important in the local adaptation to local environments, and they might be under selection driven by the environmental variables. Locus *Oke_Tf-278* (Transferrin; Elfstrom *et al*. 2007) was shown to be an outlier by both independent approaches we implemented, and its allele frequency was positively correlated with the number of frost-free days (NFFD). *Oke_Tf-278* was also shown to be under positive selection at the 95% confidence level by Seeb *et al*. (2011) in their study undertaken throughout the species’ range of *O. keta*. Evidence for positive selection at the transferrin gene among salmonid species was also found in other studies (Ford 2001). Transferrins are iron-binding proteins that are involved in iron storage and resistance to bacterial disease. Since iron is often a limiting source of nutrient in bacterial growth, transferrin may provide resistance to bacterial infection by iron binding (Ford 2001). Therefore, iron competition with salmonid pathogens could potentially act as a selective pressure on the transferrin gene. It has been found in some salmon populations that a specific transferrin genotype can confer resistance to bacterial infection (Ford 2001). Warmer temperatures have been shown to increase the infection rate of fish pathogens (Richter and Kolmes 2005). In salmonids species, populations that are already stressed by high water temperatures, manifested by weight loss, disease and displacement by better-adapted species, are more susceptible to pathogen infections (Richter and Kolmes 2005). As a result, a positive correlation between the number of frost-free days (NFFD) and the allele frequency at *Oke_Tf-278* may be explained by the presence of different intensities of bacterial infections in different environments with varying temperatures, with more infections in the south due to warmer temperatures.

*Oke_RFC2-168* was shown to be under positive selection and to correlate with two environmental variables: the number of frost-free days (NFFD) and the temperature difference between the mean warmest month temperature (°C) and the mean coldest month temperature (°C) (TD). *Oke_RFC2-168* was also shown to be under positive selection at the 99% confidence level by Seeb *et al*. (2011). *Oke_RFC2-168* is situated in the gene that codes for replication factor (*RFC*), a five-subunit DNA polymerase accessory protein, which is involved in both DNA replication and repair (Seeb *et al*. 2011). *RFC* is a protein complex that functions to facilitate the assembly of active replication complexes (Bowman et al. 2004). *RFC* is essentially a clamp loader for the sliding clamp, PCNA (proliferating cell nuclear antigen), and it can form a stable ATP-dependent complex with the sliding clamp. The a-helices and the amino acid residues that are crucial for DNA interaction are conserved in all five *RFC* subunits (Bowman et al. 2004). The SNP locus *Oke_*RFC2-168 is identified as a synonymous substitution within the coding region of the RFC2 gene (Seeb *et al*. 2011).

Overall, we observed a strong effect of environmental variables on the allele frequencies at locus *Oke_RFC2-168* (Fig. 4 and 5). An adaptation along environmental gradients, shown by a high allele frequency at *Oke_RFC2-168* where NFFD is low and TD is high (characteristic of more northern, continental climates) and a low allele frequency where NFFD is high and TD is low (characteristic of more southern and coastal climates), was also observed. The above findings suggest that the population that has a higher allele frequency at *Oke_RFC2-168* could have an adaptive advantage in extreme weather conditions where TD is high and NFFD is low. Likewise, the population that has a lower allele frequency at *Oke_RFC2-168* might be best adapted to a temperate climate where TD is low and NFFD is high. Several studies have suggested an important role of temperature and photoperiod in determining salmonid migration pattern and the physiological changes during smolt development (Skyes and Shrimpton 2010; Sykes *et al*. 2009). Increases in photoperiod and temperature can stimulate physiological changes associated with increased saltwater tolerance; temperature, along with turbidity and flow, triggers actual migration. The timing and duration of the migration of salmonid smolts are crucial to their survival in marine environment (Skyes and Shrimpton 2010; Sykes *et al*. 2009). As a result, the effect of temperature and photoperiod on salmonid smolting can lend support to important adaptive roles of both environmental variables, TD and NFFD. Yet, the SNP *Oke_RFC2-168* was identified as a synonymous substitution within the coding region of *RFC* gene, so we argue that *Oke_RFC2-168* could be linked to a favorable allele at a closely linked SNP locus, which was not examined in our study, in the coding region of the *RFC* gene.

*Oke_MARCKS-362* was found to be under positive selection and is associated with the gene that encodes myristoylated alanine-rich protein kinase C substrate (Elfstrom *et al*. 2007). The gene product of *Oke_MARCKS-362* is myristoylated alanine-rich C kinase substrate (MARCKS), a substrate for protein kinase C and is thought to be involved in cell motility, phagocytosis, membrane trafficking and mitogenesis (Dulong *et al*. 2004). We show that allele frequency at *Oke_MARCKS-362* correlates significantly with longitude. The correlation between the allele frequency at *Oke_MARCKS-362* and longitude is not straightforward and could be due to other confounding effects as we only took into considerations the abiotic factors. Biotic factors such as predation risk, female choice and pathogens can also act as selective agents in salmonid populations (Garcia de Leaniz *et al*. 2007). Therefore, biotic factors could have imposed selective pressures on chum salmon populations inhabiting along a longitudinal gradient.

Ultimately, statistical correlations between environmental variation and allele frequency differentiation do not constitute infallible evidence for natural selection (Schoville *et al*. 2012). To establish the cause and effect relationships between environmental variables and genetic variation, it is necessary to link genetic variation to phenotypic variation as well as gene functions to fitness differences (Barrett and Hoekstra 2011). This can be achieved by conducting direct measurements in the field or controlled laboratory experiments (Schoville *et al*. 2012). Moreover, functional SNPs that cause intraspecific phenotypic variation should help generate insights into the relationship between genotype and phenotype and hence the adaptive consequences of these SNPs (Macdonald and Long 2005). Further genetic sequencing and genome annotation will be required to identify functional genetic variation across genomes and to examine the adaptive functions of SNPs (Seeb *et al*. 2011; Macdonald and Long 2005).

As a concluding remark, the study of adaptive genetic variation in response to environmental variables may help predict how chum salmon populations will adapt to climate change. One prediction states that in the face of climate warming, gene migration from the pre-adapted populations in warmer climates will help to promote adaptation of the populations at the leading edge of the migrating front by bringing them better adapted alleles (Davis and Shaw 2001). Since populations at the trailing edge receive no gene flow from the pre-adapted populations, they are more likely to become locally extinct when facing the negative effects of the climate change (Davis and Shaw 2001).

## Acknowledgements

We would like to thank J. VanderKraan and B. Murray for providing helpful comments to a previous version of the manuscript.

## References

Barrett, R. D. and H. E. Hoekstra. 2011. Molecular spandrels: tests of adaptation at the genetic level. Nat. Rev. Genet. 12:767–780.

Beacham, T. D. and C. B. Murray. 1987. Adaptive variation in Body size, age, morphology, egg size, and developmental biology of chum salmon (*Oncorhynchus keta*) in British Columbia. J. Fish. Aquat. Sci. 44(2):244–261.

Bowman, G. D., M. O’Donnell and J. Kuriyan. 2004. Structural analysis of a eukaryotic sliding DNA clamp-clamp loader complex. Nature 429:724–730.

Coop, G., D. Witonsky, A. Di Rienzo, and J. K. Pritchard. 2010. Using environmental correlations to identify loci underlying local adaptation. Genetics 185:1411–1423.

Davis, M. B. and R. G. Shaw. 2001. Range shifts and adaptive responses to quaternary climate change. Science 292:673–679.

Dulong, S., S. Goudenege, K. Vuillier-Devillers, S. Manenti, P. Sylvie and P. Cottin. 2004. Myristoylated alanine-rish C kinase substrate (MARCKS) is involved in myoblast fusion through its regulation by protein kinase Ca and calpain proteolytic cleavage. Biochem J. 382:1015–1023.

Earl, D. A. and B. M. vonHoldt. 2012. STRUCTURE HARVESTER: a website and program for visualizing STRUCTURE output and implementing the Evanno method. Conserv. Genet. Resour. 4: 359–361

Eckert, A. J., A. D. Bower, S. C. Gonzalez-Martinez, J. L. Wegrzyn, G. Coop and D. B. Neale. 2010. Back to nature: ecological genomics of loblolly pine (*Pinus taeda*, Pinaceae). Mol Ecol. 19(17):3789–805.

Elfstrom C. M., C. T. Smith and L. W. Seeb. 2007. Thirty-eight single nucleotide polymorphism markers for high-throughput genotyping of chum salmon. Mol. Ecol. Notes 7:1211–1215.

Exoffier, L., T. Hofer, and M. Foll. 2009. Detecting loci under selection in a hierarchically structured population. Heredity 103:285–298.

Ford, M. J. 2001 Molecular evolution of transferrin: evidence for positive selection in salmonids. Mol. Biol. Evol. 18(4):639–47.

Garcia de Leaniz, C., I. A. Fleming, S. Einum, E. Verspoor, W. C. Jordan, S. Consuegra, N. Aubin-North, D. Lajus, B. H. Letcher, A. F. Youngson, J. H. Webb, L. A. Vøllestad, B. Villanueva, A. Ferguson and T. P. Quinn. 2007. A critical review of adaptive genetic variation in Atlantic salmon: implications for conservation. Biol Rev Camb Philos Soc. 82(2):173–211.

Limborg, M. T., S. M. Blankenship, S. F. Young, F. M. Utter, L. W. Seeb, M. H. Hansen and J. E. Seeb. 2012. Signatures of natural selection among lineages and habitats in *Oncorhynchus mykiss*. Ecol. Evol. 2(1):1–18.

Macdonald, S. J. and A. D. Long. 2005. Prospects for identifying functional variation across the genome. Proc. Natl. Acad. Sci. U S A. 102 suppl 1:6614–21.

Manel, S., B. N. Poncet, P. Legendre, F. Gugerli and R. Holderegger. 2010. Common factors drive adaptive genetic variation at different spatial scales in *Arabis alpine*. Mol. Ecol. 19:3824–3835.

McColl, G. and S.W. McKechnie. 1999. The Drosophila heat shock hsr-omega gene: an allele frequency cline detected by quantitative PCR. Mol. Biol. Evol. 16(11):1568–1574.

Pritchard, J. K., M. Stephens and P. Donnelly. 2000. Inference of population structure using multilocus genotype data. Genetics 155:945–959.

Quinn, T. P. 2005. The behavior and ecology of Pacific salmon and trout. Maryland: American Fisheries Society.

Richter, A. and A. K. Steven. 2005. Maximum temperature limits for Chinook, Coho, and Chum Salmon, and Steelhead Trout in the Pacific Northwest. Reviews in Fisheries science 13:23–49.

Sato, S., H. Kojima, J. Ando, H. Ando, R. L. Wilmot, L.W. Seeb, V. Efremov, L. LeClair, W. Buchholz, D. Jin, S. Urawa, M. Kaeriyama, A. Urano, and S. Abe. 2004. Genetic population structure of chum salmon in the Pacific Rim inferred from mitochondrial DNA sequence variation. Environ. Biol. Fish. 69:38–50.

Schoville, S. D., A. Bonin, O. Francois, S. Lobreaux, M. Christelle and M. Stephanie. 2012. Adaptive genetic variation on the landscape: methods and cases. Annu. Rev. Ecol. Evol. Syst. 43:23–43.

Seeb, L.W., W. D. Templin, S. Sato, S. Abe, K. Warheit, J. Y. Park and J. E. Seeb. 2011. Single nucleotide polymorphisms across a species’ range: implications for conservation studies of Pacific salmon. Mol. Ecol. Resour. 11:195–217.

Smith, C.T., J. Baker and L. Park et al. 2005a. Characterizaion of 13 single nucleotide polymorphism markers for chum salmon. Mol. Ecol. Notes 5:259–262.

Smith, C.T., C. M. Elfstrom, L. W. Seeb and J. E. Seeb. 2005b. Use of sequence data from rainbow trout and Atlantic salmon for SNP detection in Pacific salmon. Mol. Ecol. 14:4193–4203.

Sykes, G. E., C. J. Johnson and J. M. Shrimpton. 2009. Temperature and flow effects on migration timing of Chinook Salmon Smolts. T. Am. Fish. Soc 138:1252–1265.

Sykes, G. E. and J. M. Shrimpton. 2010. Effect of temperature and current manipulation on smolting in Chinook salmon (*Oncorhynchus tshawytscha*): the relationship between migratory behavior and physiological development. Can. J. Fish. Aquat. Sci 67:191–201.

Wang, T., A. Hamann, D. Spittlehouse and T. N. Murdock. 2012. ClimateWNA- high-resolution spatial climate data for Western North America. J. Appl. Meteorol. Clim. 61:16–29.

